# Triac treatment prevents neurodevelopmental and locomotor impairments in thyroid hormone transporter Mct8/Oatp1c1 deficient mice

**DOI:** 10.1101/2022.10.07.511125

**Authors:** Jiesi Chen, Eva Salveridou, Lutz Liebmann, Sivaraj M. Sundaram, Denica Doycheva, Boyka Markova, Christian A. Hübner, Anita Boelen, W. Edward Visser, Heike Heuer, Steffen Mayerl

**Author notes:** Equal contribution (Heike Heuer, Steffen Mayerl). **Corresponding author:** Steffen Mayerl, PhD, Dept. of Endocrinology, Diabetes & Metabolism, University Hospital Essen – Hufelandstrasse 55, D-45147 Essen – Germany, **Name and Address of corresponding authors:** Steffen Mayerl, PhD: Department of Endocrinology, University Hospital Essen, University Duisburg-Essen, Germany, Heike Heuer, Prof: Department of Endocrinology, University Hospital Essen, University Duisburg-Essen, Germany.

## Abstract

**Background:** Patients with inactive thyroid hormone (TH) transporter MCT8 display intellectual disability due to an insufficient TH transport and action in the CNS. As a therapeutic strategy, application of Triac (3, 5, 3’-triiodothyroacetic acid) and Ditpa (3, 5 -diiodo-thyropropionic acid) have been proposed as both thyromimetic compounds are not dependent on MCT8 for cellular entry. Here, we tested and directly compared the thyromimetic actions of Triac versus Ditpa in Mct8/Oatp1c1 double knockout mice (Dko), a mouse model for human MCT8 deficiency.

**Methods:** Newborn Dko mice were daily injected during the first three postnatal weeks with either Triac (50 ng/g or 400 ng/g) or Ditpa (400 ng/g or 4000 ng/g) and compared with Wt and Dko mice receiving saline injections. A second cohort of Dko mice was daily injected with Triac (400 ng/g) only between postnatal week 3 and 6. Thyromimetic effects in the CNS and peripheral tissues were monitored at different postnatal time points by immunofluorescence stainings for neural marker proteins, in situ hybridization and quantitative real time PCR. Locomotor performance was assessed in rotarod and hanging wire test. Acute brain slices of Triac treated Dko mice and their respective controls were used for electrophysiological recordings.

**Results:** Only Dko mice injected with Triac (400 ng/g) during the first three postnatal weeks showed normalized myelination, differentiation of cortical GABAergic interneurons as well as locomotor performance. Electrophysiological recordings revealed an increased frequencies of cortical spontaneous miniature inhibitory postsynaptic currents in Dko mice and a normalization of this parameter in Triac treated Dko mice. In comparison, treatment of Dko mice with Ditpa at 4000 ng/g during the first three postnatal weeks resulted in normal myelination and cerebellar development but was less effective in restoring neuronal parameters and locomotor function. Finally, Triac was more potent than Ditpa in suppressing *Trh* and *Tshb* expression, respectively, and exerts stronger thyromimetic effects in liver and kidneys.

**Conclusions:** In newborn Dko deficient mice, Triac is highly effective and more efficient than Ditpa in promoting CNS maturation and function. Yet, Triac treatment needs to be initiated directly after birth to achieve the most beneficial effects.

## Introduction

Patients carrying inactivating mutations in the thyroid hormone (TH)-specific monocarboxylate transporter 8 (MCT8) show global developmental delays and a complex cluster of severe intellectual and motor disabilities (Allan-Herndon-Dudley syndrome (AHDS)) (1–4). Additionally, patients exhibit characteristic changes in the TH serum profile with highly elevated T3 in the presence of normal/low T4 and normal/elevated TSH. According to the prevailing hypothesis, the neurological impairments of AHDS patients are caused by an insufficient TH transport into the brain while peripheral tissues rather sense the elevated serum T3 concentrations and are therefore in a hyperthyroid state (4–6). That the CNS of MCT8 patients is indeed in a TH deficient state could be substantiated by histopathological findings that include hypomyelination, decreased cerebral expression of parvalbumin (PV), a Calcium binding protein present in a distinct subset of inhibitory neurons, and a delayed cerebellar development (6, 7). Tissue-specific alterations in TH content are also a characteristic feature of Mct8 ko mice that fully replicate the abnormal serum TH profile and show increased TH concentrations in peripheral tissues. Mct8 ko mice also exhibit a diminished passage of the T3 into the CNS but are still capable of transporting T4 into the CNS (8–10). This latter observation can be explained by the presence of the T4-specific transporter Oatp1c1 in murine blood-brain barrier cells (11, 12). Indeed, only Mct8/Oatp1c1 double knock-out mice (Dko) show a strongly diminished TH content in the CNS, similar morphological alterations and disturbed neural differentiation as seen in patients, as well as profound locomotor impairments (13). In light of these similarities, Dko mice represent a suitable mouse model for AHDS and are, therefore, a valuable tool for testing therapeutic interventions.

One of the most promising therapeutic approaches is the application of TH analogs such as 3, 5-diiodo-thyropropionic acid (Ditpa) and 3, 5, 3’-triiodothyroacetic acid (Triac) that activate TH receptors and that do not depend on MCT8 for cellular entry (14–17). Studies in Mct8 deficient mice and AHDS patients have already revealed that both compounds are capable of lowering endogenous TH production and normalizing symptoms of a peripheral thyrotoxicosis (such as hypermetabolism, muscle wasting, and increased heart rate) (18–21). It is, however, still a matter of debate to which extent and during which time period both compounds can exert beneficial effects on brain maturation and function.

Here, we provide the first direct comparison of Triac versus Ditpa action on brain parameters in Dko mice. Our results clearly indicate that Triac is more effective than Ditpa in promoting a normal brain development and maturation. Moreover, our studies revealed that the most beneficial effects are obtained if Triac treatment is initiated directly after birth while a delayed treatment onset significantly compromises the efficacy of the TH analog.

## Materials and Methods

### Animals

Generation and genotyping of mice lacking concomitantly Mct8 (Slc16a2^tm1Dgen^; MGI: 3710233) and the organic anion transporting protein Oatp1c1 (Slco1c1^tm1.1Arte^; MGI: 5308446) were reported elsewhere (8, 11, 13). Heterozygous breeding pairs on C57Bl/6N background were set up to obtain Mct8/Oatp1c1 double knockout (Dko) offspring and wildtype (Wt) animals that served as controls (13). All animals were supplied with regular chow and water *ad libitum* and housed in IVC cages at a constant temperature (22°C) and light cycle (12 h light, 12 h dark). Since the Mct8 encoding gene Slc16a2 is encoded on the X-chromosome, only male mice were used in this study.

Animals were daily s.c. injected with TH analogs for indicated periods: Triac (3,5,3’-L-triioothyroacetic acid; Sigma-Aldrich) was applied at a concentration of 50 ng/g bw (T(50)) or 400 ng/g bw (T(400)) while Ditpa (3,5-diiodo-thyropropionic acid; Sigma-Aldrich) was given at a dose of 400 ng/g bw (D(400)) or 4000 ng/g bw (D(4000)). Control mice received saline injections.

For immuno-histochemical analysis, animals were intracardially perfused with 4% PFA in PBS and brains were post-fixed over-night. Coronal forebrain and sagittal cerebellar sections (50 μm) were produced with a vibratome (Thermo Scientific). Animals intended for *in situ* hybridization studies (ISH), quantitative PCR analysis (qPCR), and TH measurements were killed by CO2 inhalation. For ISH, tissues were snap-frozen in 2-methylbutane on dry-ice and stored at −80°C until further processing. Coronal forebrain cryo-sections (20 μm) and pituitary sections (14 μm) were produced with a cryostat (Leica), mounted on superfrost plus slides and stored at −80°C. For qPCR analyses, tissue was frozen on dry ice. Serum was obtained from whole blood samples collected by cardiac puncture using microvette tubes (Sarstedt) and stored at −20°C. Serum TH concentrations were determined as described elsewhere (22, 23).

### Immunohistochemistry

Staining and analysis was carried out as detailed elsewhere (13). In brief, mid-sagittal cerebellar vibratome sections were blocked and permeabilized with 10% normal goat serum in PBS containing 0.2% Triton X-100 and immunostained with a mouse anti-Calbindin D28k antibody (Sigma-Aldrich, 1:1000) followed by incubation with Alexa Fluor 555-labeled goat anti-mouse secondary antibody (Thermo Fisher, 1:1000).

Coronal forebrain sections were blocked and permeabilized as above, stained with rat anti-MBP (Millipore, 1:300), mouse anti-Parvalbumin (PV, Millipore, 1:1000) or mouse anti-GAD67 (Millipore, 1:200) and incubated with Alexa Fluor 555-labeled secondary antibody produced in goat (Thermo Fisher, all 1:1000). To visualize myelin, sections were incubated with FluoroMyelin Green Stain according to the manufacturer’s instructions (Molecular Probes, 1:300 dye dilution). Sections between Bregma 1.045 and −1.555 were employed for these analyses.

Pictures were taken with an Olympus AX70 microscope or Zeiss ApoTome2. For quantification of the Purkinje cell (PC) outgrowth, thickness of the molecular layer (ML) that reflects the dimension of the PC dendritic tree was determined at three different positions in lobules III, IV and V using ImageJ. PV positive neurons were counted in all layers of the somatosensory and retrosplenial cortex and normalized to the size of the analyzed area. For quantifying MBP, GAD67 and FluoroMyelin staining intensities, the respective integrated fluorescence signal intensities per area were measured using ImageJ software. Wt average values were set as 1.0. Blinding was achieved by attributing random numbers to the pictures. For each analysis, four brain sections per animal from 3-5 mice per experimental group were employed.

### ISH histochemistry

CDNA fragments complementary to mouse *Trh* (NM_009426.2, nt 1251-1876), mouse *Tshb* (NM_009432.2, nt 190-445), and mouse *Hr* (NM_021877.2, nt 902-1598) were used as templates for in vitro transcription using [^35^S]-UTP (Hartmann Analytik) as a substrate. Radioactively labeled cRNA probes were purified and diluted in hybridization buffer (50% formamide, 10% dextrane sulfate, 0.6 M NaCl, 10 mM Tris-HCl pH 7.5, 1× Denhardt’s solution, 100 μg/ml sonicated salmon sperm DNA, 1 mM EDTA and 0.5 mg/ml t-RNA) to a final concentration of 1 × 10^4^ cpm/μl. In order to prevent overexposure of the ISH signal, the radioactively labeled cRNA probes for *Trh* and *Tsh*b were further diluted with the respective unlabeled cRNA probes (5 ng/μl in hybridization buffer) at a ratio of 1:10 (for *Trh*) and 1:3 (for *Tshb*).

ISH was performed according to the hybridization procedures described in detail previously (13). For detection of radioactive ISH signals dehydrated sections were dipped in Kodak NTB nuclear emulsion and stored at 4°C for 3 days (*Trh* and *Tshb*) and 8 days (*Hr*). Autoradiograms were developed and analyzed under darkfield illumination with an Olympus microscope. For quantification of ISH signals, 4-6 tissue sections of 4-5 animals per experimental group were analyzed using ImageJ to determine the integrated signal intensities as described previously (13). Experiments carried out using the respective sense cRNA probes did not produce any specific ISH signals.

### qPCR

Total tissue RNA was isolated using the NucleoSpin RNA II Kit (Macherey-Nagel). cDNA synthesis was performed by reverse transcription using the Transcriptor High Fidelity cDNA-Synthesis Kit (Roche) according to the manufacture’s protocol. For each replicate, 5 ng of cDNA were employed. To exclude the presence of genomic DNA, one sample without reverse transcriptase was included as well. qPCR was performed using the iQ SYBR Green SupermixTM (Bio-Rad) and a Bio-Rad CFX384 detection system. Following primers were used:

Cyclophillin D: 5’-GCAAGGATGGCA-AGGATTGA-3’ and 5’-AGCAATTCTGCCTGGATAGC-3’; Dio1: 5’-CGTGACTCCTGAAGATGATG-3’ and 5’-CCAATGCCTATGGTTCCTAC-3’. The primer pairs were designed for an annealing temperature of 55°C. Transcript levels of Dio1 were normalized to expression levels of Cyclophillin D as a house-keeping gene. Four samples per experimental group were subjected to analysis.

### Electrophysiology

For electrophysiological recordings, 350-μm-thick brain slices were prepared from 3 weeks old mice and equilibrated in aCSF (in mM): 120 NaCl, 3 KCl, 1.3 MgSO4, 1.25 NaH2PO4, 2.5 CaCl2, 10 D-glucose, 25.0 NaHCO3, gassed with 95% O2 / 5% CO2, pH 7.3 at room temperature for at least 1 hour as described previously (24).

Patch clamp recordings were performed on coronal slices that were placed in a submerged recording chamber mounted on an upright microscope (BX51WI, Olympus). Slices were continuously superfused with gassed aCSF (2–3 ml/min, 32 °C, pH 7.3). Recordings of mIPSCs were performed using a CsCl-based intracellular solution (in mM): 122 CsCl, 8 NaCl, 0.2 MgCl2, 10 HEPES, 2 EGTA, 2 Mg-ATP, 0.5 Na-GTP, 10 QX-314 [*N*-(2,6-dimethylphenylcarbamoylmethyl) triethylammonium bromide], pH adjusted to 7.3 with CsOH. dl-APV (30 μM), CNQX (10 μM) and tetrodotoxin (0.5 μM) were added to the perfusate. mIPSCs were recorded at a holding potential of −70 mV for at least 5 min in aCSF. Data analysis was performed off-line with the detection threshold levels set to 5 pA. The following parameters were determined: frequency, peak amplitude, rise time, time constant of decay (τdecay), half-width, and electrical charge transfer.

### Behavioral studies

Motor coordination was evaluated using an Accelerating Rotarod (TSE Systems). Briefly, animals were familiarized to system by allowing them to run once with a velocity of 5 rpm. On the following day, animals were exposed to acceleration of rod rotation from 5-50 rpm within the maximum testing period of 300 seconds. The riding time of each animal was recorded twice daily and averaged for five consecutive days. Ten to fourteen animals were included in each experimental group.

Muscle strength was assessed in the hanging wire test. Animals were placed on top of a wire cage lid. After shaking gently and turning the cage upside down, the animal’s capability to cling to the wire was monitored. Time was taken until the mouse fell down or else the trial was terminated after 60 seconds. Each mouse was tested twice daily for three consecutive days and values were averaged. Twelve to sixteen mice were used per experimental group.

### Statistics

All data are presented as mean ± SD. Comparison between groups was performed by two-way ANOVA for experiments including two doses of each compound while one-way ANOVA analysis was applied for experiments using single doses followed by pairwise Tukey’s *post hoc* test, if not otherwise indicated. Differences were considered statistically significant if p < 0.05 and labelled *, p < 0.05; **, p < 0.01; ***, p <0.001; a – statistically significant difference to Wt animals; b - statistically significant difference to saline-treated Dko animals.

### Study approval

All animal procedures were in accordance with the European Union (EU) directive 2010/63/EU and approved by the Animal Welfare Committees of the Thüringer Landesamt für Lebensmittelsicherheit und Verbraucherschutz (03-011/11; TLLV; Bad Langensalza, Germany) and the Landesamt für Natur-, Umwelt- und Verbraucherschutz Nordrhein-Westfalen (84-02-04.2015.A331; LANUV; Recklinghausen, Germany).

## Results

### Comparison of Triac versus Ditpa treatment during early postnatal development

In a pilot study, we demonstrated that treatment of Dko mice between postnatal day 1 (P1) and P11 with Triac at a concentration of 50 ng/g bw (T(50)) or 400 ng/g bw (T(400)) resulted in a dose-dependent improvement of Purkinje cell (PC) dendritogenesis and myelination in the cerebral cortex at P12 (16). To directly compare the thyromimetic effects of Triac and Ditpa, we expanded this study and injected newborn Dko animals during the same postnatal period with Ditpa at either 400 ng/g bw (D(400)) or 4000 ng/g bw (D(4000).

Visualizing cerebellar PC dendritogenesis by Calbindin immunofluorescence staining confirmed a strongly reduced dendritic outgrowth in Dko mice and, consequently, a much smaller molecular layer that reached only 60% of the thickness found in Wt animals at P12 (Fig. 1A, B). Application of high doses of Triac or Ditpa to Dko mice fully restored PC outgrowth while the low dose applications were less effective. Analysis of MBP immunoreactivity in the cerebral cortex revealed similar findings. Only in Dko mice treated with Triac or Ditpa at a high dose normal MBP immunoreactivity could be achieved (Fig. 1A, C). As another indicator for thyromimetic action in the CNS, we analyzed the number of GABAergic interneurons expressing the calcium-binding protein parvalbumin (PV) in the primary somatosensory cortex as well as the retrosplenial cortex. In agreement with previous data (16), Dko mice exhibited a highly reduced number of PV+ cells in both regions (Fig. 1A, D, E). High dose Triac treatment restored 80% of the PV+ interneuron population in both areas wheras high dose Ditpa treatment improved PV+ cell numbers only to 50% of the respective Wt values.

**Fig. 1:**
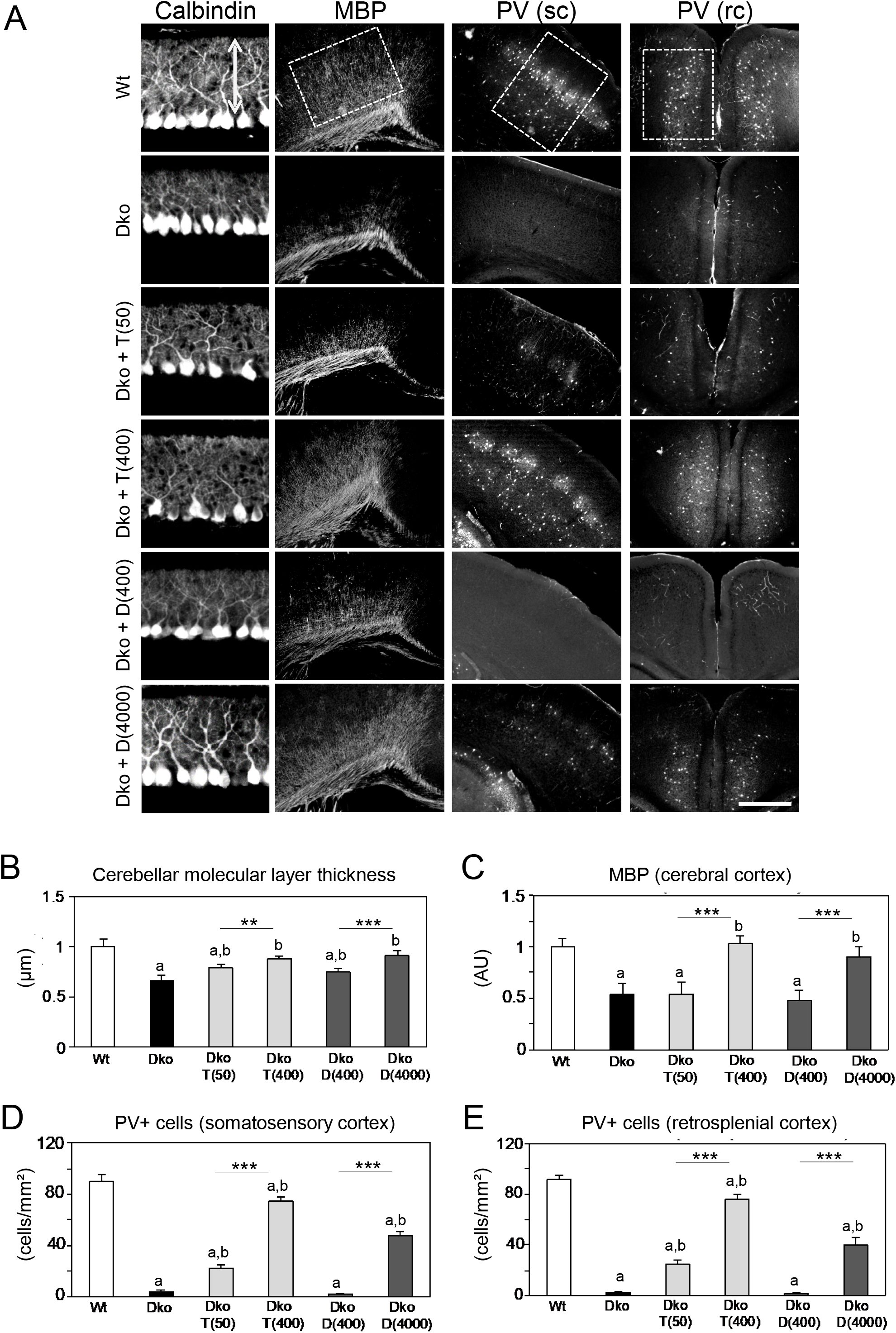
TH analog treatment initiated after birth improves brain development and maturation. Newborn mice received a daily injection of saline (control) or different doses (in ng/g bw) of TH analogs Triac or Ditpa between postnatal days 0 and 11 as indicated and brain morphology was analyzed at P12. A) Cerebellar Purkinje cell development was monitored by Calbindin immuno-staining, myelination in the cerebral cortex was assessed by MBP immuno-reactivity and PV+ interneurons were visualized in the somatosensory cortex (sc) and retrosplenial cortex (rc). B) Dimension of the Purkinje cell dendritic tree (arrow in A) was measured as a readout for the thickness of the molecular layer. Thickness was reduced in Dko animals and restored by Triac and Ditpa treatment in a dose-dependent manner. C) Integrated density of MBP-specific signals was determined in the inner layers of the cerebral cortex (box in A delineating the measured area). MBP levels were reduced in Dko animals and normalized only upon high dose Triac or Ditpa administration. Number of PV+ interneurons was enumerated in the somatosensory (D) and retrosplenial (E) cortex (boxes in A indicate areas of interest). PV+ cells were almost absent in M/O Dko mice and increased dose-dependently following Triac or high dose Ditpa application. n=3-5. Scale bars 50 μm (Calbindin), 500 μm (MBP), 250 μm (PV).

Application of TH analogs will affect the activity of the hypothalamus-pituitary-thyroid (HPT) axis by downregulating *Trh* and *Tshb* expression. To directly compare Triac versus Ditpa effects, we injected Dko mice between P1 and P20 with the same low and high doses paradigm as above and sacrificed the animals at P21. Radioactive ISH studies on brain and pituitary sections confirmed elevated *Trh* expression in hypothalamic paraventricular nucleus (PVN) neurons and increased *Tshb* expression in the anterior pituitary in Dko mice (Fig. 2A, B). Only high dose Triac treatment reduced *Trh* expression whereas both Triac concentrations caused a profound downregulation of pituitary *Tshb* mRNA expression. In response to Ditpa treatment, Dko mice only showed a mild reduction in Trh transcript expression independent of the injected dose and a moderate decrease in *Tshb* specific expression signals upon high dose Ditpa application. These data indicate that Triac exerts stronger thyromimetic action than Ditpa on the HPT axis.

**Fig. 2:**
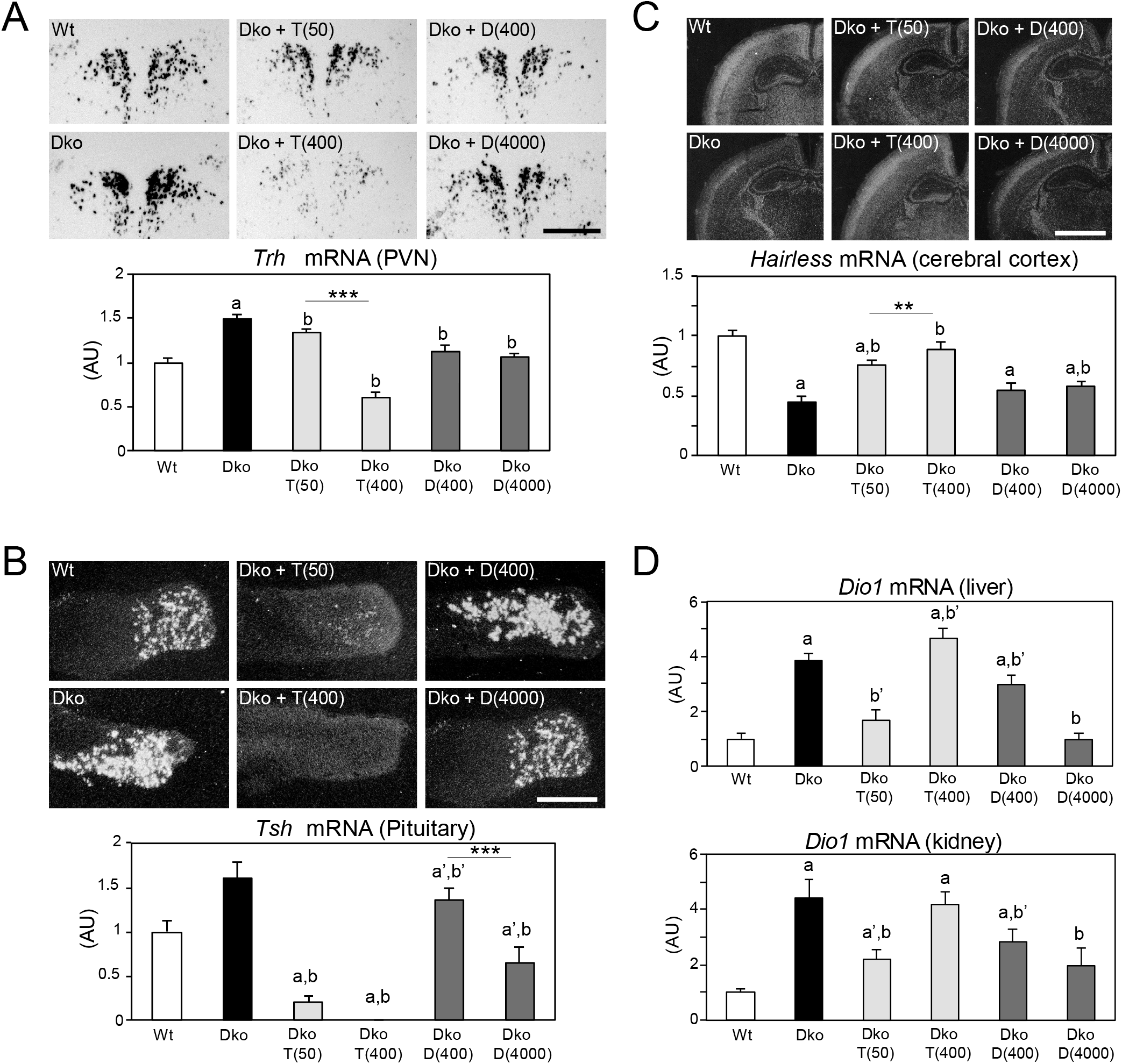
Triac executes stronger thyromimetic actions on the HPT axis than Ditpa. Dko mice were injected daily with either a saline solution or different concentrations of Triac and Ditpa between P0 and P20. At P21, activity of the HPT axis was monitored. A) Saline-treated DKo mice exhibited strongly increased Trh-specific ISH signals in PVN neurons, which were reduced in intensity following Triac application in a dose dependent manner. Ditpa had only little effect on Trh expression. Bright-field images are depicted while dark-field illuminations were employed for quantification. B) Dark-field autoradiograms illustrate alterations in Tsh beta transcripts in the pituitary with increased levels in Dko mice that were almost completely suppressed by Triac application, but only moderately so by Ditpa treatment. C) Hairless-specific hybridization signals were used to examine thyromimetic effects of TH analogs in the brain. Centrally severely hypothyroid Dko mice present with strongly decreased Hairless expression that was restored by Ditpa and, even more efficiently, by Triac application. D) Hepatic and renal Dio1 expression were studied by qPCR. Mice receiving the low dose of Triac showed a reduction in Dio1 expression in both organs while high dose Triac maintained elevated Dio1 transcript levels as seen in saline-treated Dko mice. Ditpa decreased Dio1 expression dose dependently. n=4-10. Scale bars 200 μm (TRH), 500 μm (TSH), 1 mm (Hairless).

We further assessed the expression of TH-sensitive markers in different organs to evaluate the respective tissue-specific thyroidal state. In the CNS, we analyzed mRNA expression of Hairless (*Hr*), a gene well-established to be positively regulated by TH, by ISH (Fig. 2C). *Hr*-specific signal intensities were strongly reduced in the cerebral cortex of Dko mice and increased dose-dependently following Triac treatment whereas Ditpa had only little stimulating effect on *Hr* mRNA expression in the CNS. In liver and kidney, increased *Dio1* expression (Fig. 2D) confirmed a hyperthyroid state of these tissues in Dko mice (13). Low dose Triac treatment significantly reduced *Dio1* transcript levels in both organs whereas Dko animals treated with the high dose of Triac showed similarly elevated *Dio1* levels as found in Dko mice. Low dose Ditpa treatment caused slightly reduced renal and hepatic *Dio1* expression while application of high dose of Ditpa resulted in normal hepatic *Dio1* levels and slightly elevated renal *Dio1* expression. Altogether, these results suggest that Triac exerts a much stronger thyromimetic effect than Ditpa in all analyzed organs.

### Locomotor behavior of Triac versus Ditpa treated Dko animals

We next addressed the question whether a postnatal treatment with TH analogs had any beneficial effect on locomotion. To this end, Dko mice were injected with either a high dose of Triac (400 ng/g bw) or Ditpa (4000 ng/g bw) for the first three postnatal weeks. Thereafter, TH analog treatment was terminated, and adult animals at the age of 6-7 weeks were subjected to locomotor tests.

In accordance with previous data (13), Dko mice spent significantly shorter time on the accelerating rotarod compared to control mice and did not display any visible improvement in their performance during the 5 days-training period (Fig. 3A). In contrast, Triac-treated Dko mice stayed on the rotating wheel as long as Wt mice and significantly improved their skills during the training period. The performance of Ditpa-treated Dko mice was initially not significantly different from that of Wt mice. However, these animals failed to further improve their riding time during the 5 days testing period pointing to motor learning deficits. Subsequently, animals were subjected to a hanging wire test for muscle strength assessment (Fig. 3B). Wt animals could easily cling on the metal wire for 60 s whereas Dko animals fell off after 20 s. Both Triac and Ditpa treatment significantly increased the hanging time of Dko mice.

**Fig. 3:**
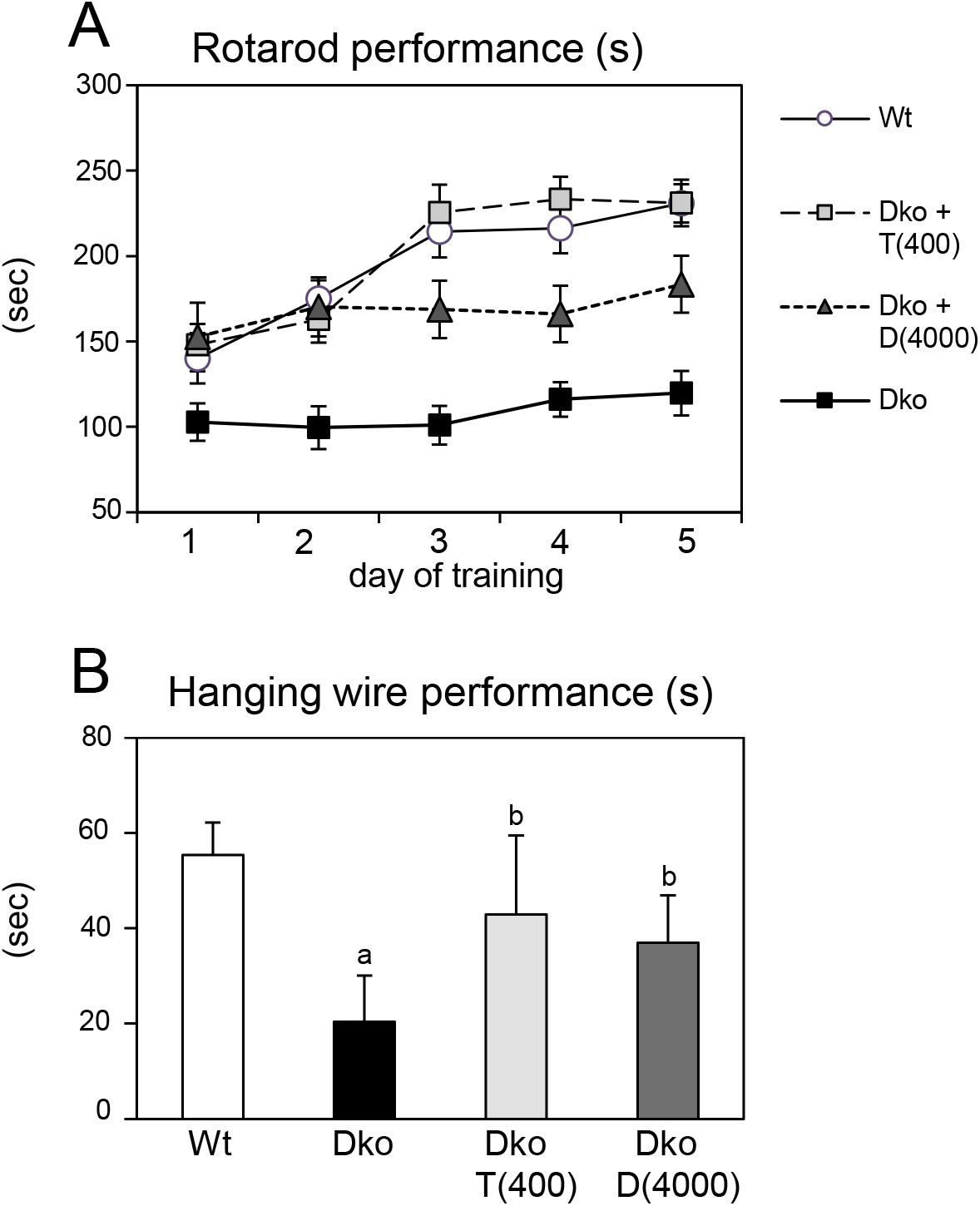
Transient early TH analog treatment improves locomotion an adulthood. Dko mice received daily injections with saline, 400 ng/g bw Triac (Dko+T(400)) or 4000 ng/g bw Ditpa (Dko+D(4000)) between P0 and P20. Thereafter, treatment was ceased and locomotor behavior was addressed at 6-7 weeks. A) In contrast to Dko mice receiving saline and showing severe locomotor deficiencies, Ditpa application moderately and Triac treatment fully normalized performance on an accelerating rotarod. B) Neuromuscular abnormalities were further examined by a hanging wire test. Dko mice clung to the wire for only a short time period, which was significantly improved in TH analog-treated experimental groups. n=10-14.

### Long-term effects of Triac versus Ditpa on the HPT axis and brain morphology

Following locomotor assessment, animals were sacrificed at the age of 10 weeks, and the activity of the HPT axis was evaluated by analyzing *Trh* and *Tshb* transcript levels by ISH and determining serum TH values. At this time point, animals have not been injected with TH analogs for 7 weeks, and, therefore, a reconstitution of the abnormal HPT axis parameters in all Dko animals could be envisioned. Hypothalamic *Trh* mRNA expression as assessed by ISH (Suppl. Fig. 1A) was equally high and serum T4 levels were equally low (Suppl. Fig. 1B) in all Dko mice independent of their treatment during the first three postnatal weeks. Serum T3 concentrations were elevated in Ditpa- and Triac-treated Dko mice, but to a significantly lesser extent than in Dko animals (Suppl. Fig. 1B). Likewise, TH-analog treated Dko animals showed lower *Tshb* expression than Dko mice (Suppl. Fig. 1A). These findings suggest that early postnatal treatment with TH analogs induces long-lasting changes in the set point of the HPT axis in Dko animals.

In addition, we investigated the long-term effects of early postnatal TH-analog treatment on brain morphology. To examine CNS myelination, coronal vibratome sections were subjected to FluoroMyelin staining and fluorescence intensities were quantified in the corpus callosum area (Fig. 4A). Compared to control animals, Dko mice showed strongly reduced myelin content. Triac and Ditpa-treated Dko mice exhibited similar FluoroMyelin staining intensities as Wt animals suggesting a normalization of myelin formation. In order to assess the maturation state of PV-expressing GABAergic neurons in retrosplenial and somatosensory cortex, PV+ cells were visualized by immunofluorescence staining. In contrast to Dko mice, Triac treated Dko animals showed similar numbers of PV+ cells in both areas as found in Wt mice (Fig. 4B). Interestingly, Ditpa treatment only slightly increased PV+ cell numbers in the somatosensory cortex, but did not exert any beneficial effect in the retrosplenial cortex. As PV is present only in a subset of GABAergic neurons, we additionally investigated expression of glutamate decarboxylase GAD67 as a key marker for all GABA producing interneurons. Quantification of GAD67 immunoreactivity in the somatosensory and retrosplenial cortex showed significantly reduced values in saline-treated Dko mice (Fig. 4C). Importantly, application of Triac, but not Ditpa restored normal GAD67 expression in Dko mice. Altogether, our data demonstrate that a transient treatment of Dko mice with TH analogs during early postnatal stages induces long lasting morphological changes in the CNS with Triac being more effective than Ditpa in normalizing brain parameters of Dko animals.

**Fig. 4:**
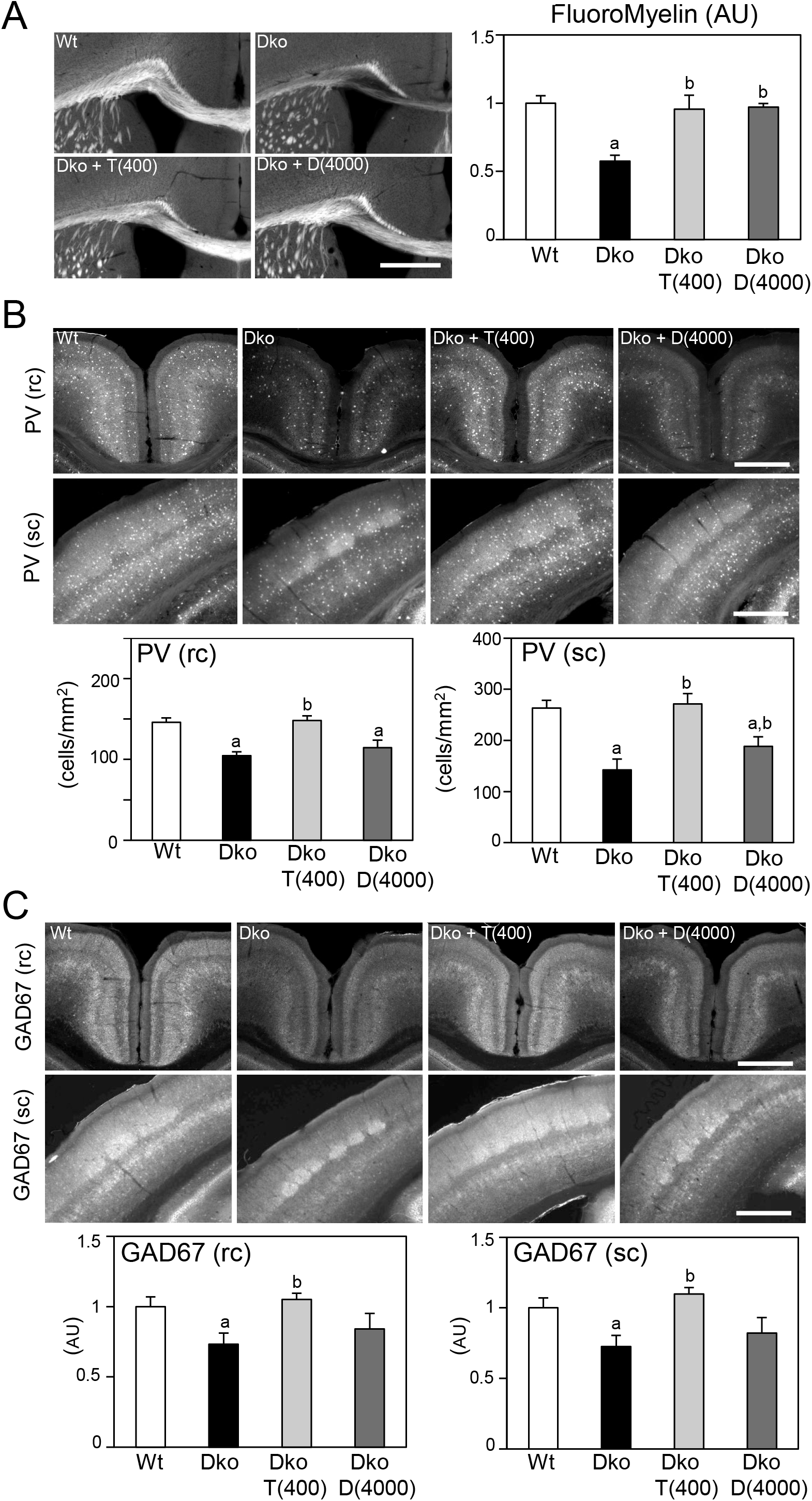
Long-term morphological alterations are induced by early postnatal TH analog application. Following a transient treatment with TH analogs Triac and Ditpa (concentrations in ng/g bw as indicated) between P0 and P20, brain parameters were examined at 10 weeks of age. A) FluoroMyelin staining was used to evaluate myelination in the corpus callosum. Hypomyelination phenotype of Dko mice could be rescued by TH analog treatment. B) PV+ interneurons were visualized in the retrosplenial cortex (rc) and somatosensory cortex (sc). Cell numbers were enumerated showing strong reductions in saline-injected Dko animals. Triac application fully restored PV+ numbers in both areas, while Ditpa only mildly improved PV+ numbers in the sc. C) GAD67 immuno-reactivity was analyzed in the same cortical regions demonstrating reduced integrated densities in saline-injected Dko mice that was unaffected by Ditpa, but fully restored upon Triac application early in life. n=3-4. Scale bars 500 μm (FluoroMyelin), 250 μm (PV, GAD67).

### Electrophysiological studies

Low GAD67 expression may result in diminished local GABA production and inhibition in the cerebral cortex that in turn will greatly influence cortical network activity. Therefore, we assessed GABAergic synaptic transmission in the somatosensory cortex in slices obtained from Wt, untreated Dko and Dko mice treated with high dose Triac for the first three postnatal weeks. Patch-clamp recordings were performed and mIPSC (spontaneous miniature inhibitory postsynaptic current) kinetics were analyzed. Representative traces recorded from pyramidal neurons in cortical layers II/III are shown in Fig. 5A. No differences in passive membrane properties (capacity: Wt 43.7±4.3 pF, Dko 36.8±1.8 pF; Dko + T(400) 35.8±2.0 pF and input resistance: Wt 67.0±4.6 MΩ; Dko 69.3±6.4 MΩ; Dko + T(400) 65.2±6.8 MΩ) were observed. In contrast, mean frequencies of mIPSCs recorded from pyramidal neurons of the somatosensory cortex were significantly increased in Dko slices compared to Wt slices (Fig.5B) suggesting that Mct8/Oatp1c1 inactivation has a major effect on GABAergic transmission in the cerebral cortex. Other recorded mIPSC kinetics parameters such as amplitudes, rise time, half-width, time constant of decay, and transported electric charges did not differ between the genotypes (Fig. 5C). Interestingly, Triac-treated Dko mice showed fully normalized mIPSC frequencies. This observation underscores the prominent and beneficial effects of Triac treatment on the development and function of the inhibitory system in Dko mice.

**Fig. 5:**
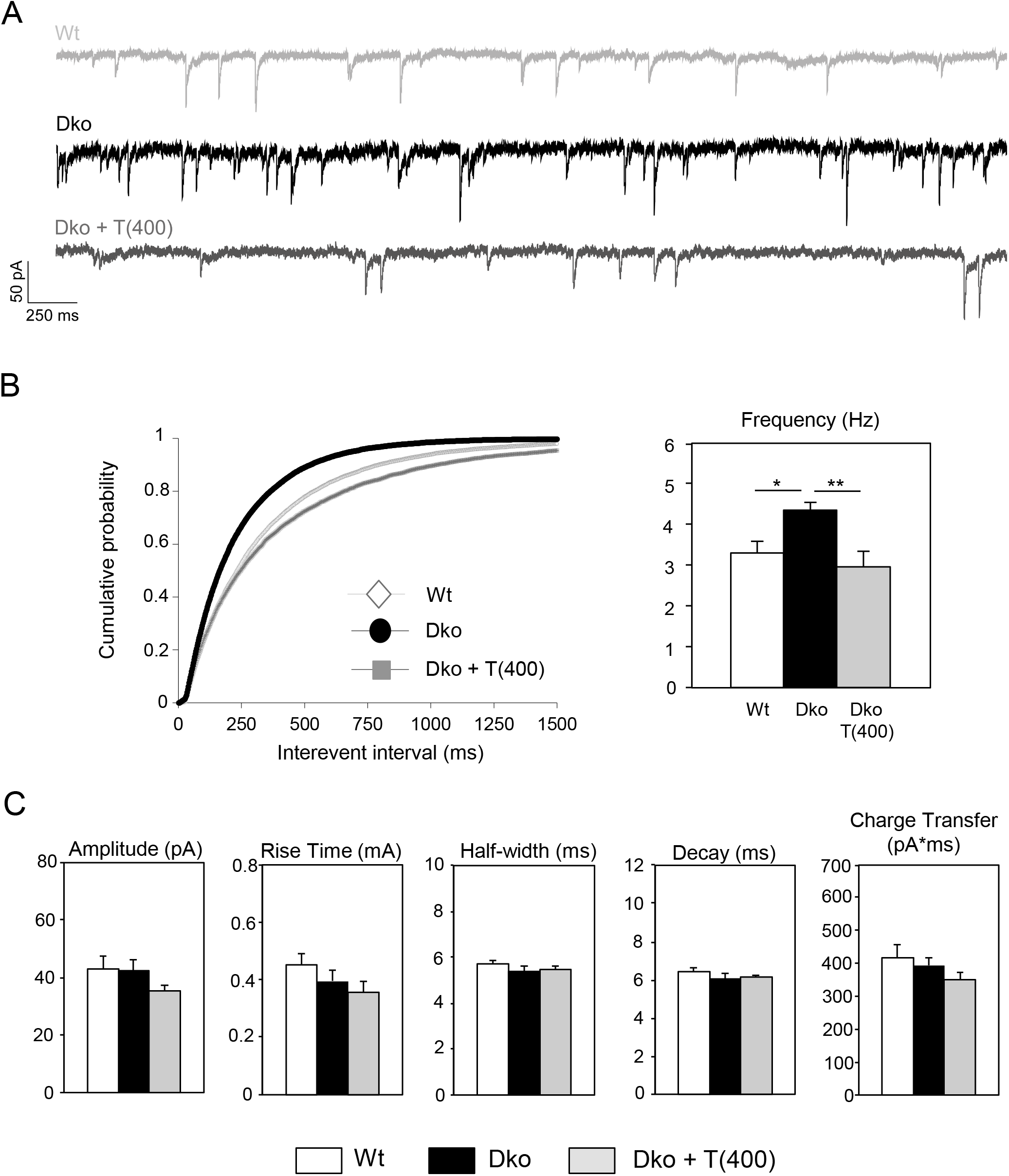
Deletion of Mct8/Oatp1c1 modulates GABAergic transmission in the cortex. Electrophysiological parameters were recorded on coronal forebrain sections using Patch-clamp technique. A) Representative traces of mIPSCs of Wt, Dko and Dko + T(400) recorded from pyramidal neurons of the somatosensory cortex. B) Cumulative plot and bar chart of mIPSC frequencies demonstrating a significant increase in Dko animals that can be rescued by application of Triac for 3 weeks. C) Amplitude, rise time, half-width, time constant of decay, and transported electric charges are not modified in neurons from Dko animals. n=17-18.

### Determining the critical time window of Triac action

Obviously, high dose Triac application restored myelination and improved maturation and function of the GABAergic system when applied between P1 and P21. In order to determine the critical time window of Triac action, we repeated our studies with a second cohort of Dko mice that only received Triac treatment between postnatal days P22-P42 (Fig. 6). Only Dko mice that received Triac during the first three postnatal weeks showed a full normalization of PV+ cell numbers in retrosplenial (Fig. 6B) and somatosensory cortex (Fig. 6C) as well as normal myelination in the corpus callosum assessed by FluoroMyelin (Fig. 6D). In contrast, in Dko mice that received Triac only between postnatal week 3 and 6, myelination was not improved while a partial recovery of PV+ immunoreactivity could still be detected. Altogether, our data indicate that the most pronounced beneficial effects of Triac on different brain parameters of Dko mice can only be achieved if postnatal treatment is initiated as early as possible.

**Fig. 6:**
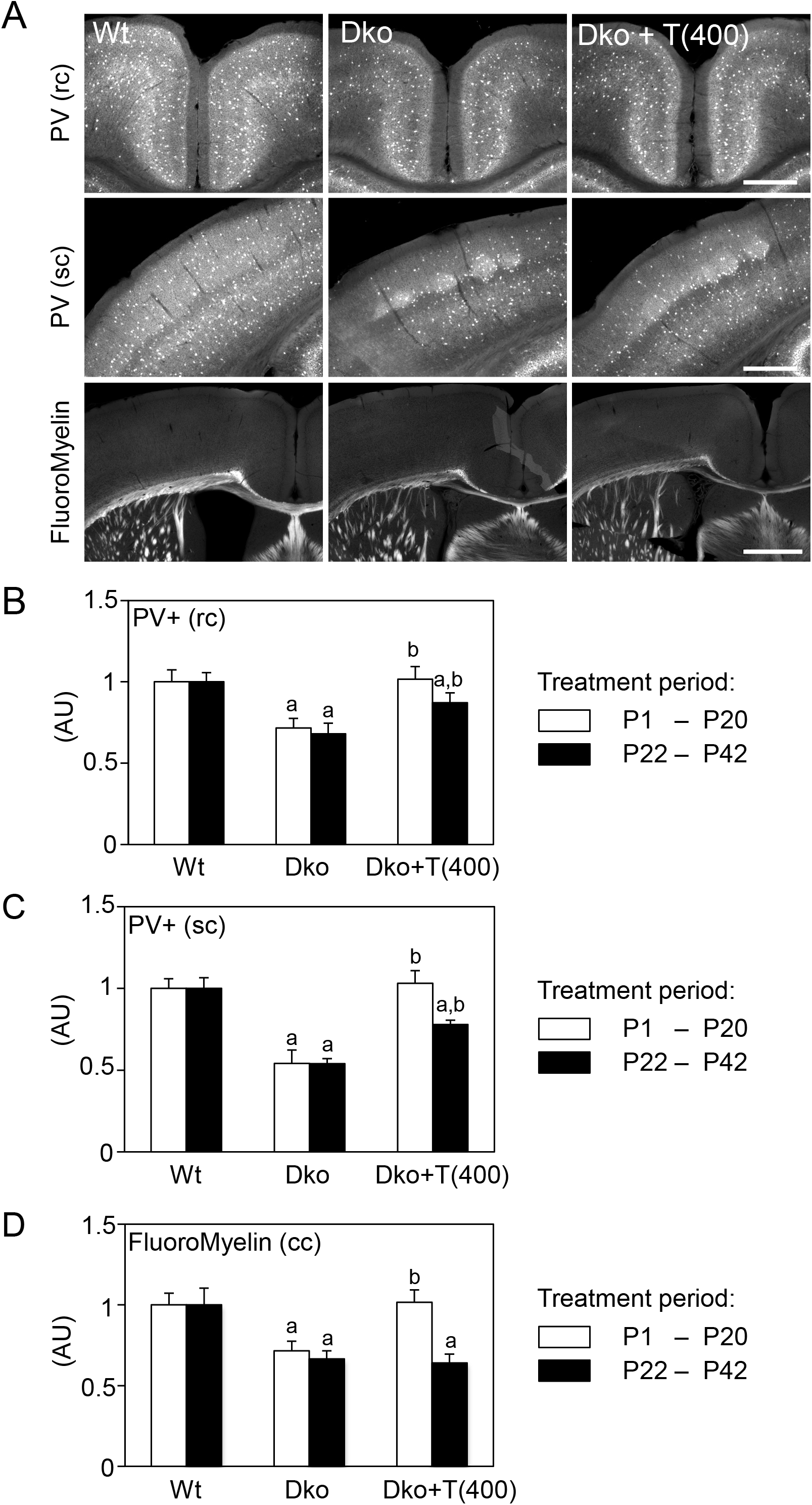
Delayed onset of Triac application compromises efficacy of the treatment. Dko mice were injected with saline or 400 ng/g bw Triac either between P0 and P20 or P22-P42 and analyzed at P43. A) PV+ interneurons were visualized in the retrosplenial cortex (rc) or somatosensory cortex (sc) and myelination was assessed in the corpus callosum by FluoroMyelin staining. Pictures of animals treated with Triac between P22-P42 are shown. B) PV+ cell numbers were quantified in the rc (B) and sc (C) and were normalized to respective Wt numbers. Low numbers of PV+ neurons in saline-injected Dko mice were fully restored following early onset Triac treatment while upon late onset only a partial recovery was observed. D) FluoroMyelin integrated density was measured in the corpus callosum (cc). Hypomyelination phenotype of Dko mice could only be rescued by Triac application between P0-P20. Late onset of the treatment did not have any effects. n=3-4. Scale bars 250 μm (PV), 500 μm (FluoroMyelin).

## Discussion

AHDS represents a devastating neurodevelopmental disease for which no therapy is currently registered. Early treatment strategies such as the application of T4 in combination with PTU in order to block T4 to T3 conversion aimed to normalize serum TH parameters and to ameliorate the symptoms of peripheral thyrotoxicosis (25). Such a treatment regiment, however, will not improve neurological symptoms given that a diminished TH transport across brain barriers and thus a reduced TH action inside the CNS represent the major pathogenic mechanisms. This uptake blockage may be circumvented by applying thyromimetic substances that bypass MCT8 and are able to activate TH receptors in neural target cells. In a compassionate study, Ditpa was given for 26-40 months to four affected children at the age of 8-25 months and was able to both normalize high serum T3 concentrations and reduce symptoms of hypermetabolism (18). Improvements in neurological functions, however, have not been reported. Likewise, in a first international clinical trial with forty-six AHDS patients (median age 7.1 years) which were treated with Triac for 12 months, a rapid reduction of the high serum T3 levels as well as beneficial and sustainable effects regarding clinical sings of a peripheral thyrotoxicosis could be observed (21). This Triac trial was not designed to detect any impact of the treatment on neurodevelopmental outcomes in human MCT8 deficiency although a trend towards neurodevelopmental improvement was noted in a subset of patients in an exploratory analysis. Thus, it is still an open question whether Triac (and Ditpa) treatment has any long-lasting beneficial effects on CNS development. A second clinical phase 2 study, Triac trial II (NCT02396459), has recently been initiated to specifically assess Triac effects in patients younger than 30 months of age at the start of the treatment whereas a similar clinical trial for Ditpa is in preparation (NCT04143295).

Although both TH analogs have already been tested in different model systems (15, 16, 19, 26–30), previous studies did not allow to draw any conclusions as to which of the two compounds exerts stronger beneficial effects on CNS development due to a variety of confounding factors (i.e. usage of different Mct8 mutants, treatment regimens, read-out parameters and drug concentrations). The major aim of our study was therefore to evaluate and directly compare the thyromimetic potential of Ditpa and Triac in Mct8/Oatp1c1 double knock-out (Dko) mice as a preclinical AHDS model. Due to a profound CNS specific TH deprivation, these Dko mice exhibit distinct brain histomorphological abnormalities (hypomyelination, compromised cerebral and cerebellar neuronal differentiation) that were also seen in AHDS patients (6, 7, 13). Moreover, by detecting a rise in cortical mIPSC frequencies in Dko mice (Fig. 5), we provide another relevant read-out parameter that underscores the impact of M/O deficiency on inhibitory neuronal network activity.

Here, we provide a first systematical testing of different concentrations of Triac and Ditpa in parallel. Two doses from previous concentration-finding pilot studies were employed for Triac (16). For Ditpa, we used 10-fold higher concentrations compared to Triac as such an order-of-magnitude-increase was needed to achieve similar thyromimetic effects in cerebellar Purkinje cell outgrowth *in vitro* (Kersseboom; not published). Likewise, 10-times higher concentration of Ditpa compared to Triac were needed to induce myelin marker p0 mRNA expression in mct8 mutant zebrafish to a similar extent (28). Moreover, we focused on brain parameters as read-outs that are well-established TH targets and that have been reported to be affected in AHDS patients as well as in our AHDS mouse model of Mct8/Oatp1c1 double deficiency (7, 13).

As major findings of our study, we could demonstrate for the first time robust and long-lasting beneficial effects of Triac on various brain parameters in Dko mice only if a high-dose Triac treatment was initiated directly after birth. This regimen was sufficient to fully restore cerebellar development, myelination as well as maturation of cortical GABAergic neurons in Dko mice as evidenced by immunofluorescence analysis (Fig. 1). Cortical mIPSC frequencies recorded in acute brain slices and elevated in Dko animals, showed normal values in high-dose Triac treated Dko animals (Fig. 5). Possibly, these electrophysiological findings reflect the altered expression pattern of PV and other Calcium binding proteins in Dko mice as a reduced neuronal Ca^2+^ buffering capacity will cause an elevated release probability and thus a rise in mIPSC frequency. Finally, high dose Triac treatment of Dko mice restored locomotor performance as assessed by rotarod and hanging wire test (Fig. 3). In comparison, high dose Ditpa treatment during the first three postnatal weeks also stimulated myelination and cerebellar development but was significantly less effective in mending cortical GABAergic neuron maturation and normalizing locomotor function, suggesting that Triac has a stronger potential for the treatment of MCT8 patients.

Several aspects, however, have to be considered before translating these preclinical data into clinical practice. The high concentration of Triac needed for effective normalization of brain parameters in our mouse model resulted in a fully downregulated HPT axis with *Tshb* transcript below the detection limit (Fig. 2A,B). Although we were not able to determine serum T riac versus TH concentrations in Triac treated animals due to antibody cross-reactivity issues (31) we speculate that endogenous serum TH concentrations are highly reduced in Triac treated Dko mice. Triac is most likely the only TH receptor active substance in these animals with such a high thyromimetic activity (at the highest dose) that it still causes a peripheral thyrotoxic state as indicated by the elevated hepatic and renal *Dio1* expression (Fig. 2D). This scenario is certainly in contrast to the situation of the AHDS patients in the first clinical Triac trial (21). In fact, one major aim of this clinical study was to achieve a reduction in the highly elevated serum T3 levels in order to ameliorate peripheral thyrotoxicosis while T4 serum levels should still be detectable. To that end, an average dose of 38.3 μg/kg/day was given, which is in the same range as the low Triac dose (T(50)) used in this study. However, treating Dko mice with such a low Triac dose had only little beneficial effects on brain parameters such as myelination or interneuron maturation (Fig. 1). It is therefore tempting to speculate that with respect to the neurological outcome, MCT8 patients might benefit more from a treatment with a higher dose of Triac even if under these circumstances signs of e.g. hepatic thyrotoxicosis are still present.

Another intriguing observation is the optimal time window of Triac application. In our study, only the early onset Triac treatment regiments exerted strong beneficial effects on brain parameters in Dko while late onset Triac application starting at postnatal day 22 and lasting for 3 weeks did not improve myelination and only insufficiently restored cortical PV+ cell numbers (Fig. 6). Why the CNS of Dko mice showed very little response to the late-onset Triac treatment is still elusive. As one hypothesis, the not-yet identified transport systems by which Triac enters the CNS and neural target cells might be down-regulated in the mature mouse CNS (26). Additionally, the critical time window during which neural differentiation processes are still sensitive to TH might be closed. The latter scenario would underscore the utmost importance of an early postnatal diagnosis of AHDS followed by an immediate treatment initiation.

As an alternative to TH analog treatment, gene therapy approaches exploiting AAV vector constructs have been considered (32–34). These interventions aim to express a functional MCT8 transporter in brain endothelial cells and eventually restore TH action in other neural cell types. Intravenous injection of endothelial cell specific AAV-BR1-Mct8 constructs in newborn Dko mice resulted in improved cerebellar development, myelination and GABAergic marker expression (34). Yet, these beneficial effects were less profound compared to the alterations seen here upon high dose Triac treatment in Dko mice. Treatment of juvenile Dko mice with AAV-BR1-Mct8 at P30 induced expression of well-established T3-target genes, yet failed to improve myelination and only slightly induced GABAergic marker expression, similar to our observations with late onset Triac treatment. Likewise, in Dko mice treated at P30 with AAV9-MCT8 constructs, serum TH parameters remained abnormal indicating that a peripheral hyperthyroidism is preserved, while TH-target genes in the CNS showed only a partial response (33).

By comparing different treatment strategies and similar read-out parameters in the same Dko mouse model, Triac appears as the most promising treatment approach for AHDS to date. Yet, Triac presumably needs to be given at a high dose and the treatment should be initiated as early as possible to achieve the best outcome for the patients.

## Author contributions

JC, SM, WEV and HH devised the study. JC, ES, SMS, DD and SM conducted the in vivo work and all histo-morphological analyses. LL performed and analysed the electrophysiological experiments. AB determined serum TH levels. JC, LL, CAH, AB, SM and HH interpreted the results. JC, SM, WEV and HH wrote the manuscript.

## Acknowledgments

This work was supported by the BMBF within the E-RARE project “THYRONVERVE” (01GM1401), by grants of the Deutsche Forschungsgemeinschaft to HH (HE3478/7-1; 8-1 as well as CRC/TR 296; P01; P09), to SM (CRC/TR 296; P19) and by Sherman family funds to HH. We would like to thank Markus Korkowski for excellent technical assistance.

---

**Suppl. Fig. 1:**
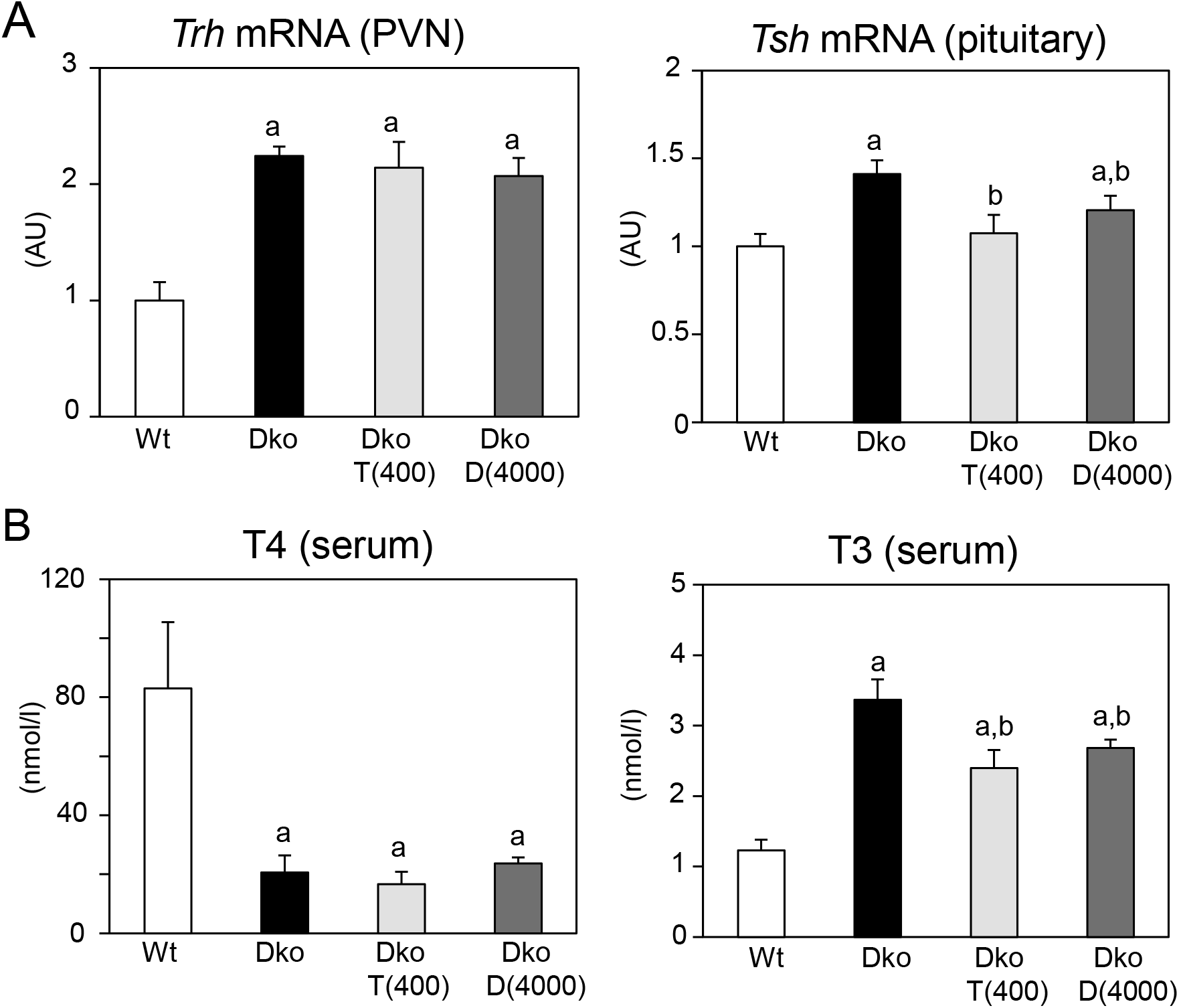
Early postnatal TH analog application permanently modulates the set point of the HPT axis. Activity of the HPT axis was monitored at 10 weeks of age in animals receiving TH analogs Triac or Ditpa daily between P0 and P20 only. A) Radioactive ISH was performed and Trh mRNA expression in the PVN and levels of Tsh beta transcripts in the pituitary were quantified. Following cessation of TH analog treatment, Trh levels returned to abnormally high values as seen in saline-injected Dko mice. In the pituitary, Tsh expression was still decreased in animals that received an early, transient treatment with TH analogs. B) TH serum levels were determined. T4 was equally low in all Dko animals independent of the treatment, whereas T3 concentrations significantly reduced in Triac and Ditpa-treated Dko mice in comparison to saline-injected genotype controls. n=3-5.

